# WHIX is a T6SS secretion domain found in polymorphic double-edged sword effectors

**DOI:** 10.1101/2024.08.21.608442

**Authors:** Chaya Mushka Fridman, Kinga Keppel, Vladislav Rudenko, Jon Altuna-Alvarez, David Albesa-Jové, Eran Bosis, Dor Salomon

## Abstract

Gram-negative bacteria employ the type VI secretion system (T6SS) to deliver toxic effectors into neighboring cells and outcompete rivals. Although many effectors have been identified, their secretion mechanism often remains unknown. Here, we describe WHIX, a domain that is sufficient to mediate the secretion of effectors via the T6SS. Remarkably, we find WHIX in T6SS effectors that contain a single toxic domain, as well as in effectors that contain two distinct toxic domains fused to either side of WHIX. We demonstrate that the latter, which we name double-edged sword effectors, require two cognate immunity proteins to antagonize their toxicity. Furthermore, we show that WHIX can be used as a chassis for T6SS-mediated secretion of multiple domains. Our findings reveal a new class of polymorphic T6SS cargo effectors with a unique secretion domain that can deploy two toxic domains in one shot, possibly reducing recipients’ ability to defend themselves.

## Introduction

Interbacterial competition can be a strong driver of bacterial evolution [1,2]. A major player in bacterial warfare is the type VI secretion system (T6SS), an offensive apparatus widespread in gram-negative bacteria [3–7]. The T6SS delivers a cocktail of antibacterial toxins, called effectors, into neighboring bacteria in a contact-dependent manner [5,8–12]. Upon T6SS activation, a contractile sheath propels a tube that is made of hexameric Hcp rings and capped with a spike comprising VgrG, and PAAR or PAAR-like repeat-containing proteins (the latter will be collectively referred to as PAAR hereafter) out of the cell; the tube-spike, which is decorated with effectors, then penetrates a neighboring bacterium where the effectors are deployed [13].

Many T6SS effector families have been investigated to date, revealing toxic mechanisms in the bacterial cytoplasm, periplasm, and membrane, including peptidoglycan-degrading enzymes [8,14,15], pore-forming toxins [16–18], phospholipases [10,19], nucleases [20–23], NAD(P)^+^-degrading enzymes [24,25], effectors inhibiting protein synthesis [26] or cell division [27], and ADP-ribosyl transferases [27,28]. Importantly, antibacterial effectors are encoded next to a cognate immunity protein that prevents self/kin-intoxication [5,29–31].

T6SS effector repertoires include “specialized effectors”—tube-spike components fused to a toxic domain, and “cargo effectors”—toxic domain-containing proteins that non-covalently bind a secreted tube-spike component [32,33], often via a “loading platform” region at its C-terminus [34–36]. Some cargo effectors require an adaptor protein for proper loading [37,38], or a secreted adaptor or co-effector for secretion [39,40].

Cargo effectors that load onto the spike tend to be modular proteins comprising a C-terminal toxic domain and an N-terminal secretion domain responsible for loading the effector onto the spike “loading platform” [21,40–43]. We previously identified several non-structural T6SS secretion domains named MIX [41], RIX [40], and PIX [43], each defining a class of polymorphic T6SS effectors. We showed that these domains are necessary and sufficient to mediate secretion through the T6SS. An additional domain, FIX, was also used to define a widespread class of polymorphic T6SS effectors, yet its role in T6SS secretion remains unclear [23].

Although many T6SS effectors have been described, we hypothesize that additional T6SS secretion domains, which can be used to reveal new classes of polymorphic T6SS effectors, are yet to be identified.

Here, we describe a T6SS secretion domain named WHIX. We reveal that WHIX domains define a widespread class of polymorphic effectors containing periplasm-targeting toxic domains. Remarkably, we demonstrate that WHIX domains, which are sufficient to mediate T6SS-depended secretion, can be fused to either one or two toxic domains, each requiring its own cognate immunity protein for protection.

## Results

### WHIX is a widespread T6SS-specific domain found in peptidoglycan-targeting effectors

In previous work, we determined the effector repertoires of T6SS1 in *Vibrio alginolyticus* 12G01 [44] and *V. coralliilyticus* BAA-450 [45]. While examining these effector repertoires, we observed that two proteins, one from *Vibrio alginolyticus* (WP_005373349.1) and the other from *V. coralliilyticus* (WP_006961879.1), share a highly similar N-terminus (65% identity) and a less similar C-terminus (37% identity). Furthermore, their downstream-encoded immunity proteins are not similar (Fig 1A). This observation led us to hypothesize that the highly similar N-termini of these effectors contain a previously undescribed T6SS secretion domain.

**Fig 1.**
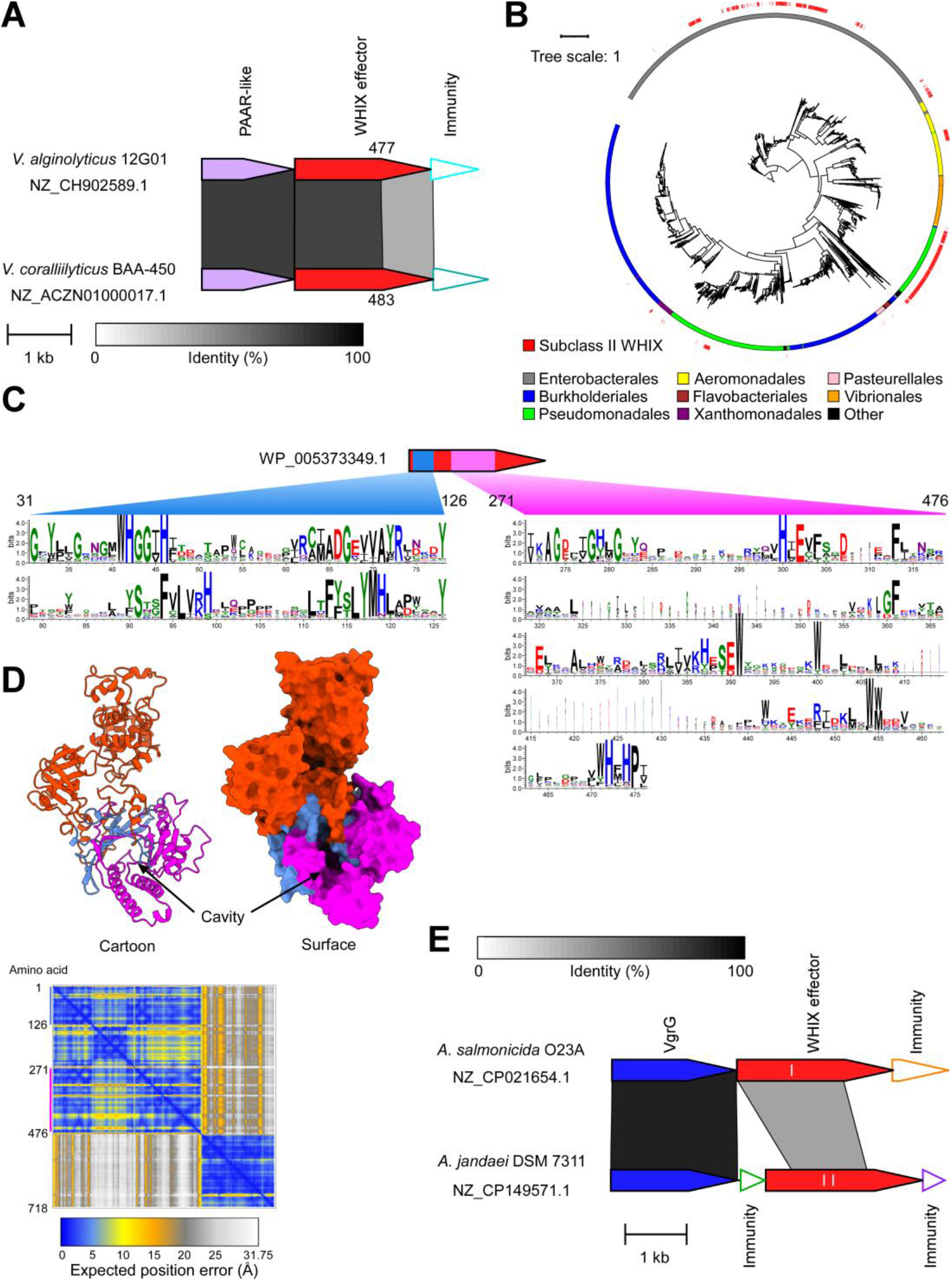
WHIX domain defines a widespread family of T6SS effectors. **(A)** Gene synteny of the T6SS effectors WP_005373349.1 from *V. alginolyticus* and WP_006961879.1 from *V. coralliilyticus*. Gray rectangles denote amino acid identity between regions. Bacterial strain and Genbank accession are denoted. **(B)** Phylogenetic distribution of WHIX domain-containing proteins. The bacterial order and the WHIX subclass are denoted by color. The evolutionary history was inferred by using the Maximum Likelihood method and JTT+I+G4 model. The tree is drawn to scale, with branch lengths measured in the number of substitutions per site. Evolutionary analyses were conducted using IQ-TREE. **(C)** Conservation logo based on a multiple sequence alignment of WHIX domain sequences. The position numbers correspond to the amino acids in WP_005373349.1. **(D)** An AlphaFold 3 structure prediction of WP_005373349.1, as ribbon and surface representations. The first region of WHIX is colored blue; the second region is colored magenta. The AlphaFold Predicted Aligned Error (PAE) plot is shown below. The color key represents the expected position error for each pair of residues in Å units. Blue represents low predicted errors, indicating high confidence in the relative positions of those residues. The orange color indicates higher predicted errors, suggesting lower confidence. **(E)** Gene synteny of subclass I and II WHIX effectors, WP_087755869.1 from *A. salmonicida* O23A and WP_082035413.1 from *A. jandeai* DSM 7311, respectively. Gray rectangles denote amino acid identity between regions. Bacterial strain and Genbank accession are denoted.

Following these observations, we set out to identify sequences homologous to the N-terminus of the *V. alginolyticus* effector in other bacterial proteins. Computational analyses revealed 4,434 unique protein accession numbers encoded by 12,071 genomic loci spread across various gram-negative bacterial orders, such as Enterobacterales, Burkholderiales, Pseudomonadales, and Aeromonadales (Fig 1B and Dataset S1). Notably, >98% of the genomes encoding these homologs contain a T6SS (Dataset S2).

A multiple sequence alignment of the homologous sequences revealed a bipartite conserved motif. The first region corresponds to amino acids 31-126 and the second corresponds to amino acids 271-476 in the *V. alginolyticus* effector (Fig 1C). We named this domain WHIX after a conserved WHxxxH motif found in the first region (<u>WH</u> type s<u>IX</u> motif). The predicted structure of the *V. alginolyticus* effector, generated with AlphaFold 3 [46], suggests that the bipartite WHIX domain forms a distinct folded module with a deep cavity (Fig 1D and File S1). The predicted C-terminal toxic domain appears to form a separate module.

Analysis of WHIX-containing proteins revealed two subclasses (Fig 1B and Fig 1E). In subclass I (e.g., WP_087755869.1), the WHIX domain is located at the N-terminus whereas in subclass II (∼16% of the unique protein accession numbers; e.g., WP_082035413.1), an additional extension, > 99 amino acids long, is found N-terminal to WHIX; the latter appear to reside predominantly in Enterobacterales, Pseudomonadales, and Aeromonadales genome (Fig 1B).

The majority of WHIX-containing proteins (96.5% of the unique protein accession numbers) contain a C-terminal extension. When examining the identity of these C-terminal extensions, we found diverse domains predicted to target the peptidoglycan. These include, for example, domains belonging to the Glyco_hydro_19 superfamily (cl46694), lyz_endolysin_autolysin (cd00737), NLPC_P60 superfamily (cl21534), and Muramidase superfamily (cl13324) (Dataset S1). In instances in which WHIX-containing proteins lack a C-terminal extension, they can be found upstream of genes encoding a predicted peptidoglycan-degrading enzyme (e.g., WP_227130466.1 and WP_223384031.1). Analyses of the N-terminal extensions found in subclass II WHIX-containing proteins suggested that they also comprise diverse peptidoglycan-targeting enzymes (Fig S1, Table S1, and File S2); in two instances, WHIX is fused to an N-terminal VgrG domain. Further inspection of the genes immediately downstream and upstream of the predicted toxic domains fused to WHIX revealed that they predominantly encode proteins with a predicted signal peptide (Dataset S1), indicating possible periplasmic localization.

Moreover, some of these proteins contain domains previously implicated in toxin antagonism and T6SS immunity, such as the DUF1311/LrpI lysozyme inhibitor [47,48] and DUF3828/Tai3 [49] (Dataset S1), suggesting that the genes flanking the WHIX-encoding gene encode the cognate immunity protein to the adjacently-encoded toxic domain. Taken together, the above observations led us to conclude that WHIX defines a new class of polymorphic antibacterial T6SS effectors.

### The Aeromonas jandaei T6SS is an active antibacterial system

To further investigate WHIX effectors, we characterized the T6SS in *Aeromonas jandaei* DSM 7311 (also named CECT 4228 and ATCC 49568), which is predicted to encode a subclass II WHIX effector (Dataset S1). Since the publicly available genome of this strain was not assembled into a chromosome (NCBI RefSeq assembly GCF_000819955.1), we re-sequenced the genome, resulting in a complete chromosome assembly (NCBI Reference Sequence: NZ_CP149571.1). Analysis of the newly assembled genome revealed a main T6SS gene cluster with a predicted Rhs-containing effector similar to the previously reported TseI [50], and two auxiliary modules containing T6SS tube-spike components: one encoding a Tle1-like effector [51] and the other a subclass II WHIX effector, WP_082035413.1, which we named Awe1 (*<u>A</u>eromonas* <u>W</u>HIX <u>e</u>ffector <u>1</u>) (Fig 2A).

**Fig 2.**
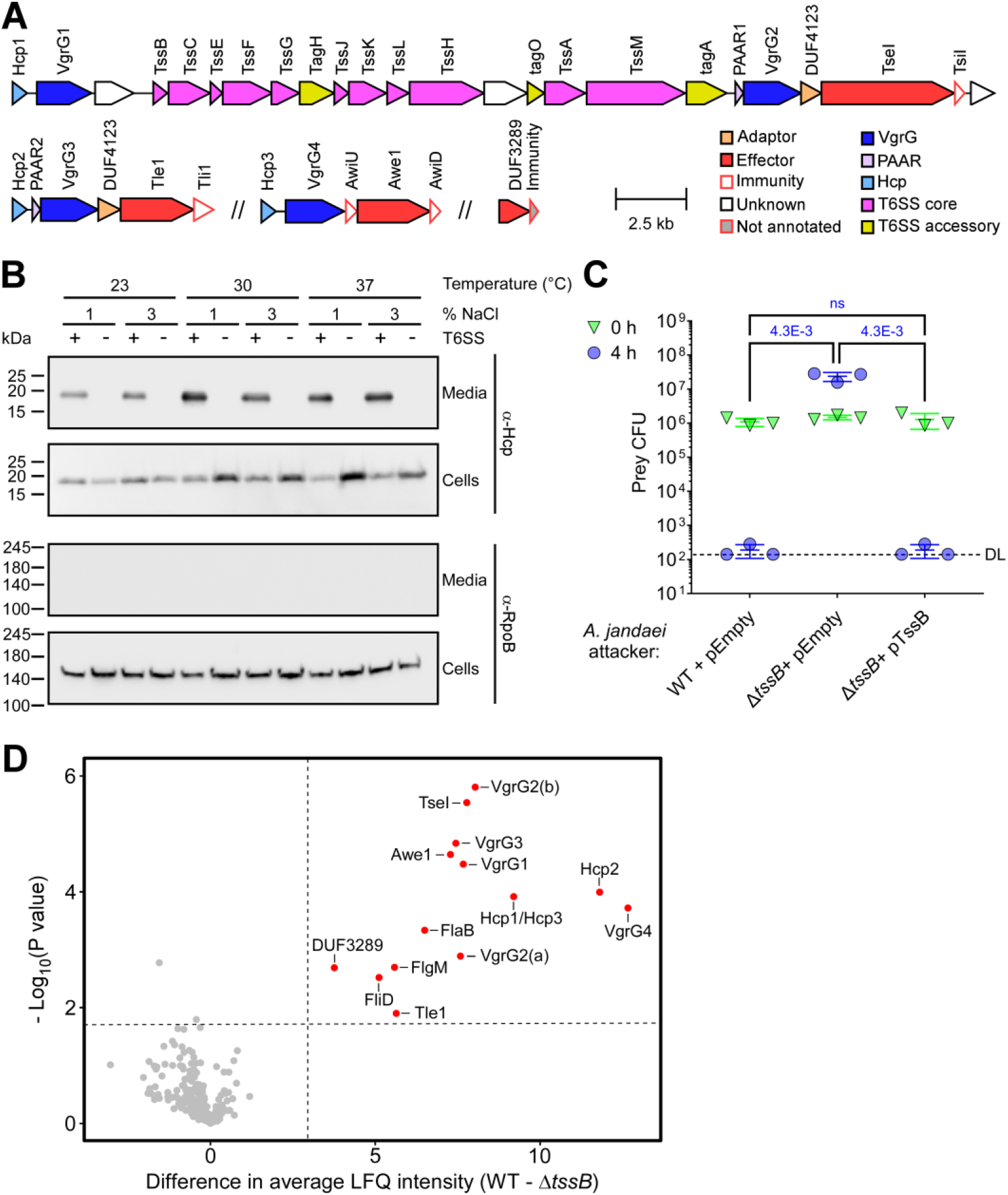
*A. jandaei* DSM 7311 has an antibacterial T6SS that secretes four effectors. **(A)** Schematic representation of *A. jandaei* DSM 7311 T6SS gene cluster and auxiliary operons. Predicted protein activity is denoted above. **(B)** Expression (cells) and secretion (media) of Hcp from wild-type (WT; T6SS^+^) *A. jandaei* DSM 7311 and a T6SS^−^ mutant strain (Δ*tssB*) grown for 3 hours at the indicated temperatures in media containing 1% or 3% (wt/vol) NaCl (LB or MLB, respectively). RNA polymerase beta subunit (RpoB) was used as a loading and lysis control. Results from a representative experiment out of at least three independent experiments are shown. **(C)** Viability counts (colony forming units; CFU) of *E. coli* MG1655 prey strains before (0 h) and after (4 h) co-incubation with the indicated *A. jandaei* DSM 7311 attacker strains containing an empty plasmid (pEmpty) or a plasmid for the arabinose inducible expression of tssB (pTssB), on LB plates supplemented with 0.1% (wt/vol) L-arabinose at 30°C. The statistical significance between samples at the 4 h time point was calculated using an unpaired, two-tailed Student’s *t* test; ns, no significant difference (*P* > 0.05); WT, wild-type; DL, the assay’s detection limit. Data are shown as the mean ± SD; *n* = 3. The data shown are a representative experiment out of at least three independent experiments. **(D)** Volcano plot summarizing the comparative analysis of proteins identified in the media of *A. jandaei* DSM 7311 WT and T6SS^−^ (Δ*tssB*) strains, using label-free quantification (LFQ). The average LFQ signal intensity difference between the WT and T6SS^−^ strains is plotted against the -Log_10_ of Student’s *t* test *P* values (*n* = 3 biological replicates). Proteins that were significantly more abundant in the secretome of the WT strain (difference in average LFQ intensities > 3; *P* value < 0.02; with a score > 40) are denoted in red.

To identify the conditions in which this T6SS is active, we monitored the expression and secretion of the hallmark secreted T6SS component, Hcp [52], using a custom-made antibody designed to detect all three Hcp proteins encoded by *A. jandaei* (Fig 2A). As shown in Fig 2B, the T6SS is active between 23°C and 37°C, in media containing either 1% or 3% (wt/vol) NaCl (LB or MLB media, respectively). Hcp secretion was T6SS-dependent, since it was not observed in a T6SS^−^ strain in which we deleted the core component *tssB*. Secretion of Hcp was restored when *tssB* was complemented from a plasmid (Fig S2). Because the highest level of Hcp secretion was observed when bacteria were grown in LB media (1% [wt/vol] NaCl) at 30°C, we used these conditions for subsequent assays.

The three effectors that we computationally identified in *A. jandaei* (i.e., TseI, Tle1, and Awe1) are encoded next to putative immunity genes (Fig 2A). Therefore, we hypothesized that this T6SS plays a role in interbacterial competition. To test this hypothesis, we used the wild-type *A. jandaei* and its T6SS^−^ mutant strain (Δ*tssB*) as attackers in competition against *Escherichia coli* MG1655 prey. Both wild-type and the *tssB*-complemented mutant attacker strains killed the *E. coli* prey during a 4-hour co-incubation on LB agar plates at 30°C, as evidenced by a drop in prey viability; the T6SS^−^ mutant did not (Fig 2C). Taken together, these results indicate that the T6SS in *A. jandaei* is active under a wide range of growth conditions in which it mediates antibacterial toxicity.

### Aeromonas jandaei T6SS secretes four antibacterial effectors

To characterize the T6SS effector repertoire of *A. jandaei*, we next employed a comparative proteomics approach. Using mass spectrometry, we compared the proteins secreted by the wild-type *A. jandaei* (T6SS^+^) with those secreted by a Δ*tssB* mutant strain (T6SS^−^) when grown in LB media at 30°C. We identified 13 proteins that were significantly enriched in the supernatant of the wild-type strain (Fig 2D and File S3). These include the four structural VgrG proteins (note that VgrG2 is separated into ‘a’ and ‘b’ because the mass spectrometry data were analyzed against a genome annotation in which the gene is split into two open reading frames; File S3), the three structural Hcp proteins (note that Hcp1 and Hcp3 are identical and are thus annotated together), and the three computationally predicted effectors (TseI, Tle1, and Awe1). In addition, we identified three flagella proteins (FlaB, FlgM, and FliD); notably, manipulation of the T6SS was previously reported to affect motility in other *Aeromonas* species [53,54]. In addition, we identified a protein containing DUF3289, WP_042032936.1.

Because DUF3289 was recently proposed to be an antibacterial toxic domain that targets the bacterial periplasm [40,55,56], we hypothesized that the protein identified in our comparative proteomics analysis, WP_042032936.1, is a T6SS effector. In agreement with the comparative proteomics results, WP_042032936.1 was secreted in a T6SS-dependent manner when a tagged version of the protein was expressed from a plasmid in *A. jandaei* (Fig S3A). Although not annotated in the NCBI RefSeq genome, we identified a short open reading frame immediately downstream of the DUF3289-encoding gene (between position 2,059,343 and 2,059,057 in genome NZ_CP149571.1), encoding a predicted 95 amino acid long hypothetical protein with an N-terminal signal peptide and three transmembrane helices (analyzed using Phobius [57]) (Fig 2A). We hypothesized that this is an immunity protein antagonizing the DUF3289-containing effector. Indeed, deletion of the predicted effector and immunity pair rendered a prey strain sensitive to attack by a wild-type *A. jandaei* attacker and not by a T6SS^−^ mutant strain (Δ*tssB*), as evidenced by the decrease in prey viability over time (Fig S3B). Expression of the predicted immunity gene from a plasmid restored the prey strain’s ability to antagonize the T6SS-mediated attack. These results indicate that the DUF3289-containing WP_042032936.1 and its downstream-encoded protein are a T6SS effector and immunity pair. Furthermore, taken together, our results demonstrate that the *A. jandaei* T6SS secretes at least four antibacterial effectors.

### WHIX is sufficient for T6SS-mediated secretion of multiple domains

Next, we focused our investigation on Awe1, a subclass II WHIX-effector predicted to contain two toxic domains, one on each side of WHIX (Fig 1E). Prediction of the Awe1 structure using AlphaFold 3 revealed a similar fold to that of WP_005373349.1 from *V. alginolyticus*, containing a cavity made by a bipartite WHIX domain (Fig 3A and File S4). In addition, two domains, one at the N-terminus and one at the C-terminus of Awe1, were apparent in the prediction. Furthermore, when the Awe1 structure was predicted together with the putative immunity proteins encoded upstream (Awi1U) and downstream (Awi1D) of Awe1 (Fig 2A), we found that these immunity proteins are expected to bind the N-terminal and C-terminal domains of Awe1, respectively (Fig 3B and File S5). These results imply that the N- and C-terminal domains fused to WHIX are toxic domains. The N-terminal domain (amino acids 1-147) is predicted to belong to the EnvC superfamily (according to NCBI conserved domain database; CDD [58]), which includes peptidoglycan peptidase domains [59]. In agreement, the AlphaFold 3-predicted structure of this domain is similar to the M23 peptidase domain of the ShyA endopeptidase from *V. cholerae* (PDB: 6UE4A), as determined by Foldseek analysis [60] (Fig S4A and File S6). In contrast, no similarity to known domains was identified for the Awe1 C-terminal domain (amino acids 700-862) by sequence homology. Nevertheless, the AlphaFold 3-predicted structure of this domain was found to share low similarity with the predicted structure of the FlgJ peptidoglycan hydrolase from *V. parahaemolyticus* (AlphaFold identifier: Q9X9J3), as determined by Foldseek analysis (Fig S4B and File S6). Based on these observations, we hypothesized that WHIX is a T6SS secretion domain that can carry two toxic domains, one on each of its ends.

**Fig 3.**
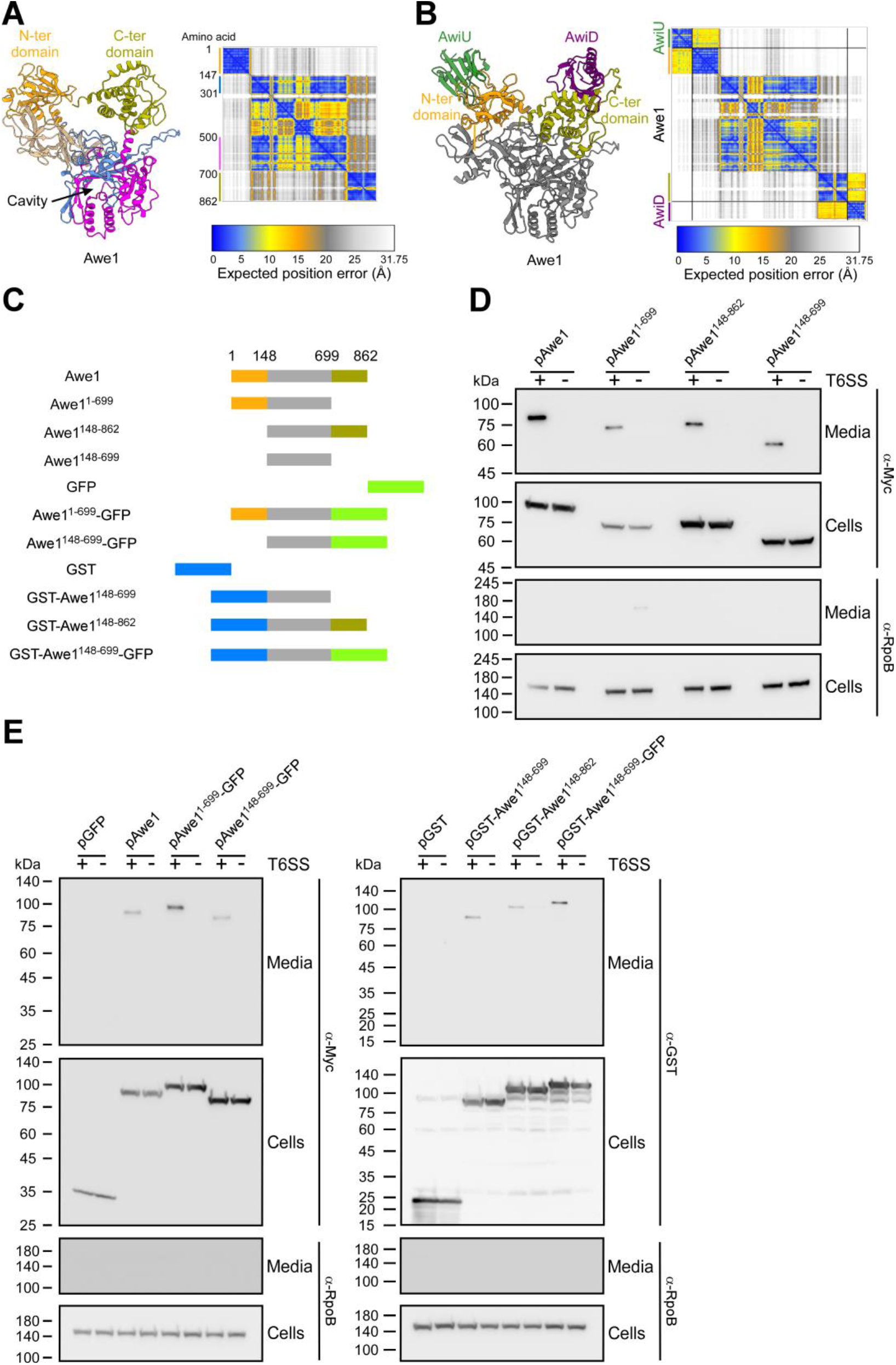
WHIX can carry two flanking domains for secretion via the T6SS. AlphaFold 3 structure predictions of Awe1 **(A)** or Awe1 in complex with AwiU and AwiD lacking their predicted N-terminal signal peptides **(B)**. The first part of the Awe1 WHIX domain is colored blue (amino acids 148-301); the second part of WHIX (amino acids 500-699) is colored magenta; the N-terminal (N-ter) Awe1 domain fused to WHIX (amino acids 1-147) is colored orange; the C-terminal (C-ter) Awe1 domain fused to WHIX (amino acids 700-862) is colored beige; AwiU is colored green; AwiD is colored purple. The AlphaFold Predicted Aligned Error (PAE) plot is also shown; black lines denote the borders between proteins included in the prediction. The color key represents the expected position error for each pair of residues in Å units. Blue represents low predicted errors, indicating high confidence in the relative positions of those residues. The orange color indicates higher predicted errors, suggesting lower confidence. **(C)** Schematic representation of the Awe1 truncations and chimeras used in the secretion assays shown in panels D-E. **(D-E)** Expression (cells) and secretion (media) of the indicated C-terminally Myc-tagged or N-terminally GST-tagged Awe1 forms expressed from an arabinose-inducible plasmid (pAwe1) in Δ*awe1 A. jandaei DSM 7311* (T6SS^+^) or Δ*awe1*/Δ*tssB* (T6SS^−^) mutant strains grown for 3 hours at 30°C in LB media supplemented with kanamycin and 0.05% L-arabinose. GFP, superfilder GFP. RNA polymerase beta subunit (RpoB) was used as a loading and lysis control. Results from a representative experiment out of at least three independent experiments are shown.

To determine whether the Awe1 WHIX domain is sufficient to mediate T6SS-dependent secretion, we monitored the secretion of the full-length Awe1 or its N- and C-terminal truncated forms (Fig 3C) from *A. jandaei* in which we deleted the genomic *awe1* (Δ*awe1*; T6SS^+^) or from a derivative in which we inactivated the T6SS (Δ*awe1*/Δ*tssB*; T6SS^−^). Truncation of the Awe1 N-terminal domain (Awe1^148-862^), C-terminal domain (Awe1^1-699^), or both domains together (Awe1^148-699^), did not hamper the T6SS-dependent secretion of Awe1 (Fig 3D), indicating that the WHIX domain alone is sufficient for T6SS secretion.

To further support WHIX’s role as a T6SS secretion domain that can carry two domains, one on each end, we monitored the secretion of Awe1 in which the N-terminal domain was swapped with a glutathione S-transferase (GST) domain and the C-terminal domain was swapped with a superfolder green fluorescent protein (GFP) (Fig 3C). As shown in Fig 3E, the Awe1 WHIX domain (amino acids 148-699) was able to carry two non-T6SS domains, fused to its N- and C-terminal ends either individually or together, for secretion via the T6SS. These results implicate WHIX as a T6SS secretion domain able to mediate the secretion of two flanking domains.

### Awe1 is a double-edged sword effector

The findings described above support the hypothesis that Awe1 contains two distinct toxic domains and is thus akin to a double-edged sword. Based on this hypothesis, we first sought to determine whether the two distinct toxic domains in Awe1 are functional. Using *E. coli* as a surrogate bacterial cell, we found that Awe1 is toxic when sent to the bacterial periplasm (by fusing it to an N-terminal PelB signal peptide) (Fig 4A and Fig S5). In support of the hypothesis that both the N- and C-terminal domains of Awe1 are functional toxic domains, we observed that expression of either is detrimental to *E. coli* when sent to the periplasm. Notably, the periplasmic expression of the WHIX domain alone is not toxic to *E. coli*, and its effect on *E. coli* growth is comparable to that of the full-length Awe1 expressed in the cytoplasm.

**Fig 4.**
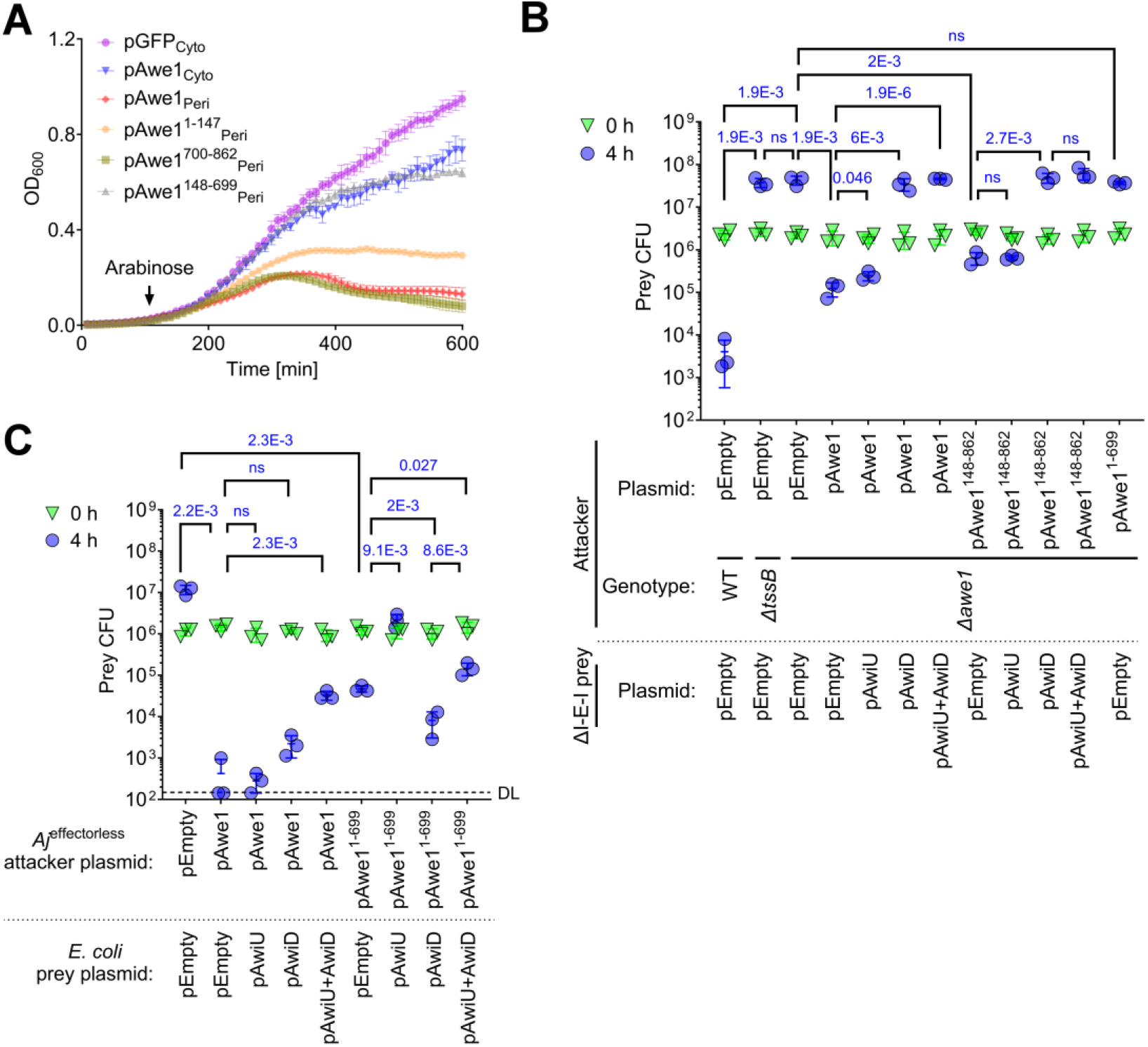
Awe1 is a “double-edged sword” effector. **(A)** Growth of *E. coli* MG1655 containing plasmids for the arabinose-inducible expression of superfolder GFP (GFP) or the indicated Awe1 forms. An arrow denotes the timepoint at which L-arabinose (0.05% [wt/vol]) was added to the LB media. Cyto, cytoplasmic expression; Peri, periplasmic expression by fusing the protein to an N-terminal PelB signal peptide. Data are shown as the mean ± SD; *n* = 4 biological samples. The experiments were repeated three times with similar results. Results from representative experiments are shown. **(B-C)** Viability counts (CFU) of *A. jandaei* DSM 7311 prey strains in which we deleted the genes encoding AwiU, Awe1, and AwiD (ΔI-E-I) (B) or of *E. coli* BL21(DE3) prey strains (C) containing an empty plasmid (pEmpty) or a plasmid for the arabinose-inducible expression of AwiU (pAwiU) or AwiD (pAwiD) before (0 h) and after (4 h) co-incubation with the indicated *A. jandaei* DSM 7311 attacker strains containing an empty plasmid or a plasmid for the arabinose-inducible expression of the indicated Awe1 form, on LB plates supplemented with 0.05% (in B) or 0.1% (in C) (wt/vol) L-arabinose at 30°C. The statistical significance between samples at the 4 h time point was calculated using an unpaired, two-tailed Student’s *t* test; ns, no significant difference (*P* > 0.05); WT, wild-type; DL, the assay’s detection limit. Data are shown as the mean ± SD; *n* = 3. The data shown are a representative experiment out of at least three independent experiments.

Next, we sought to determine whether the two Awe1 toxic domains require two distinct cognate immunity proteins (i.e., AwiU and AwiD) to antagonize them. To this end, we generated an *A. jandaei* strain in which we deleted the genes encoding the predicted effector Awe1 and both predicted immunity proteins AwiU and AwiD (ΔI-E-I), and we used it as prey in self-competition assays. This strain was killed by a wild-type *A. jandaei* attacker during competition, as evidenced by the decrease in viability over four hours of co-incubation on an LB agar plate (Fig 4B). The killing was mediated by the T6SS-delivered Awe1, since inactivation of the T6SS (Δ*tssB*) or of Awe1(Δ*awe1*) enabled prey growth during the co-incubation period. Expression of the full-length Awe1 from a plasmid in the Δ*awe1* attacker strain restored its ability to kill the sensitive prey, as did the expression of an Awe1 truncated version containing only the WHIX domain fused to the C-terminal toxic domain (Awe1^148-862^). Expression of AwiD from a plasmid in the prey restored its ability to antagonize both full-length Awe1 and its truncated version lacking the predicted N-terminal toxic domain (Awe1^148-862^). Surprisingly, however, the expression of an Awe1 truncation containing only the WHIX domain fused to the N-terminal toxic domain (Awe1^1- 699^) was not toxic towards the prey strain, and the AwiU predicted immunity protein did not antagonize Awe1-mediated toxicity during co-incubation (Fig 4B). These results suggest that although the C-terminal toxic domain of Awe1 is functional during self-competition and is antagonized by the downstream encoded AwiD, the predicted N-terminal toxic domain of Awe1 is not toxic under these conditions.

Because the N-terminal domain of Awe1 is toxic when expressed in *E. coli* (Fig 4A), we hypothesized that although it is a functional toxic domain, a yet-unknown non-immunity protein-resistance mechanism abrogates its toxicity toward *Aeromonas* in self-competition assays. A similar observation was previously made with the peptidoglycan-targeting effector TseH from *V. cholerae* [61]. Therefore, we set out to determine whether the N-terminal Awe1 domain can intoxicate an *E. coli* prey in a T6SS-dependent manner and whether AwiU is able to antagonize it. To this end, we first constructed an *A. jandaei* mutant strain in which we deleted the four known effectors (*Aj*^effectorless^) to obtain an effectorless strain unable to intoxicate *E. coli* (Fig 4C). Expression of the full-length Awe1 from a plasmid in the effectorless attacker strain restored its ability to kill *E. coli* prey. In contrast to the results obtained in the *Aeromonas* self-competition assay (Fig 4B), AwiD alone was unable to antagonize the full-length Awe1-mediated toxicity when it was expressed from a plasmid in the *E. coli* prey strain (Fig 4C). However, co-expression of both AwiU and AwiD from a plasmid significantly restored the prey strain’s ability to antagonize the attack. Furthermore, in contrast to the results obtained in the *Aeromonas* self-competition assay (Fig 4B), the expression of a truncated version of Awe1 containing only the N-terminal toxic domain and the WHIX domain (Awe1^1-699^) intoxicated the *E. coli* prey strain, and the expression of AwiU alone in the prey strain was sufficient to antagonize this toxicity (Fig 4C). Taken together, these results confirm our hypothesis that Awe1 is a double-edged sword effector containing two toxic domains, one on each end of the WHIX domain, and requiring two cognate immunity proteins to antagonize their toxicity.

### The C-terminus of a cognate VgrG is required for Awe1 delivery

We hypothesized that Awe1 is loaded onto the upstream-encoded VgrG4 (WP_198493475.1) for delivery via the T6SS. In agreement with our findings that the Awe1 WHIX domain is sufficient for T6SS-dependent secretion, an AlphaFold structure prediction suggests that the Awe1 WHIX domain is loaded onto the C-terminal tail of VgrG4 by capping it (Fig 5A and File S7).

**Fig 5.**
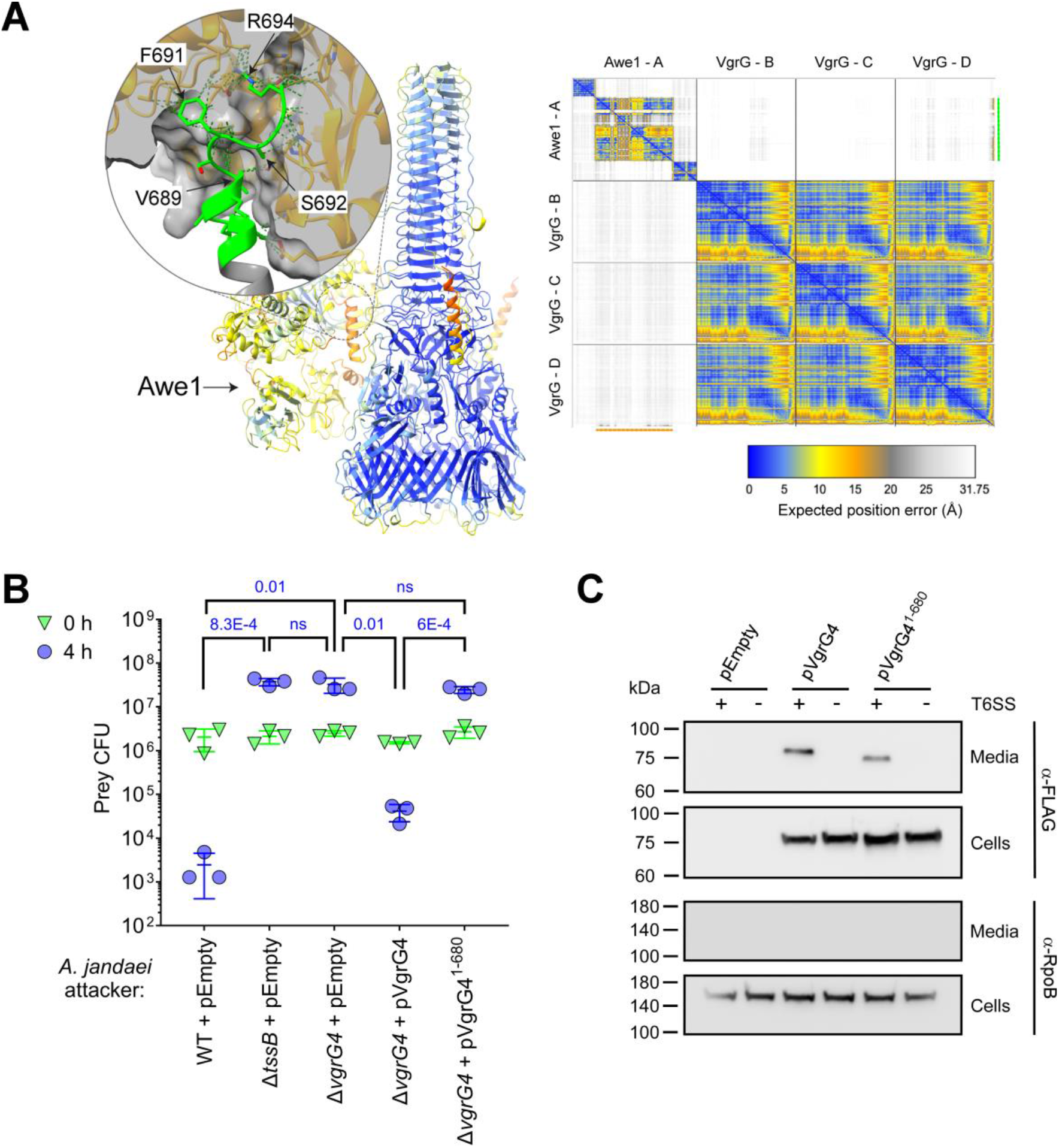
The C-terminus of VgrG4 is required for Awe1 delivery. **(A)** AlphaFold structure prediction of the complex assembled by an Awe1 monomer and a VgrG4 trimer, shown as a ribbon representation. The color gradient from blue to orange correlates with the Predicted Aligned Error (PAE) values, where blue represents regions with a very high prediction confidence level, and orange represents regions with a very low confidence level. The inset is a close-up view of the predicted Awe1-VgrG4 interacting region. VgrG4 and Awe1 residues predicted to interact with each other are represented in green and orange, respectively. Predicted interactions are shown with dotted lines. The C-terminal residues of VgrG4: Lys684, Ser685, Lys688, Val689, Ser690, Phe691, Ser692, Gly693, and Arg694 are predicted to interact with residues within the WHIX domain cavity of Awe1; for clarity, only four VgrG4 residues are labeled. The AlphaFold Predicted Aligned Error plot is shown on the right. The color key represents the expected position error for each pair of residues in Å units. Blue represents low predicted errors, indicating high confidence in the relative positions of those residues. The orange color indicates higher predicted errors, suggesting lower confidence. Awe1 (chain A) and VgrG4 (chain D) residues interacting are underscored with orange (x-axis) and green lines (y-axis), respectively. These regions correspond to the 11 C-terminal residues of VgrG4 and the WHIX domain of Awe1, respectively. **(B)** Viability counts (CFU) of *A. jandaei* DSM 7311 prey strains in which we deleted the genes encoding AwiU, Awe1, and AwiD (ΔI-E-I) before (0 h) and after (4 h) co-incubation with the indicated *A. jandaei* DSM 7311 attacker strains containing an empty plasmid (pEmpty) or a plasmid for the arabinose-inducible expression of the indicated N-terminally FLAG-tagged VgrG4 forms, on LB plates supplemented with 0.05% (wt/vol) L-arabinose at 30°C. The statistical significance between samples at the 4 h time point was calculated using an unpaired, two-tailed Student’s *t* test; ns, no significant difference (*P* > 0.05); WT, wild-type. Data are shown as the mean ± SD; *n* = 3. **(C)** Expression (cells) and secretion (media) of the N-terminally FLAG-tagged VgrG4 forms used in panel B, expressed from an arabinose-inducible plasmid (pVgrG4) in *A. jandaei* DSM 7311 Δ*vgrG4* (T6SS^+^) or a T6SS^−^ mutant strain (Δ*vgrG4*/Δ*tssB*) grown for 3 hours at 30°C in LB media supplemented with chloramphenicol and 0.005% L-arabinose. RNA polymerase beta subunit (RpoB) was used as a loading and lysis control. In B and C, results from a representative experiment out of at least three independent experiments are shown.

To determine whether the C-terminus of VgrG4 is required for Awe1 delivery, we constructed a strain in which we deleted *vgrG4*. When this Δ*vgrG4* strain was used as an attacker in a self-competition assay against a ΔI-E-I prey strain (specifically sensitive to intoxication by Awe1), it was unable to intoxicate the prey, similar to the T6SS^−^ attacker Δ*tssB* (Fig 5B). These results indicated that VgrG4 is required for Awe1 delivery. Moreover, although expression of the full-length VgrG4 from an arabinose-inducible plasmid restored antibacterial activity to the Δ*vgrG4* attacker strain, expression of VgrG4 in which we truncated the 14 C-terminal amino acids (VgrG4^1-680^) was unable to restore toxicity (Fig 5B). Notably, the truncated VgrG4 form was secreted in a T6SS-dependent manner (Fig 5C), implying that the absence of Awe1-mediated intoxication in the presence of the truncated VgrG4 resulted from the effector’s inability to properly load onto the spike. Taken together, these results indicate that Awe1 requires the C-terminus of VgrG4 for T6SS-mediated delivery into prey cells.

## Discussion

In this work, we describe a widespread class of polymorphic T6SS effectors that share a domain named WHIX. We show that the WHIX domain is sufficient to mediate secretion via the T6SS, possibly by loading onto a C-terminal tail of a secreted spike component. Importantly, we demonstrate a unique trait of WHIX domains—they are found in classic effectors containing one toxic domain, as well as in effectors akin to a double-edged sword containing two distinct toxic domains, each requiring its own cognate immunity protein.

WHIX domains are found almost exclusively in proteins encoded by bacterial genomes harboring a T6SS. Taken together with the proximity of many WHIX-encoding genes to T6SS tube-spike components (e.g., VgrG and PAAR; Dataset S1) and the fact that WHIX-containing proteins have been demonstrated to secrete via the T6SS in bacteria from different orders (i.e., Vibrionales and Aeromonadales) [44,45,62], we conclude that WHIX-containing proteins are T6SS effectors.

We show that WHIX effectors can be divided into subclass I, in which WHIX is at the N-terminus and usually fused to a C-terminal toxic domain, and subclass II, containing two toxic domains fused to either side of WHIX. Bifunctional T6SS effectors containing two catalytic domains targeting the peptidoglycan were previously described by Le *et al* [63]. However, in contrast to subclass II WHIX-effectors, these bifunctional effectors appear to require a single cognate immunity protein to antagonize their toxicity. Here, we demonstrate that subclass II WHIX effectors, akin to double-edged swords, contain two toxic domains that function separately, each requiring its own cognate immunity protein to prevent intoxication. Notably, Jiang *et al* recently described an effector with two catalytic domains in *A. veronii* [64], which is a homolog of Awe1; however, misannotation of the upstream-encoded immunity protein as a DUF4123 family adaptor, in combination with investigating its toxicity only in self-competition assays, possibly prevented its identification as a double-edged sword effector.

We hypothesize that WHIX domains’ ability to carry two fused toxic domains reduces the risk of resistance developing in prey strains, as well as expands the potential target range of a single effector. Indeed, we find that the N-terminal toxic domain of Awe1 does not intoxicate *Aeromonas* in self-competition, yet it is functional against *E. coli* prey in which both the downstream-encoded immunity protein and the upstream-encoded immunity protein are required to antagonize Awe1-mediated toxicity. Moreover, the unique ability of the WHIX domain to carry two flanking domains, either toxic or non-T6SS related, during T6SS-mediated secretion makes it a promising chassis for the future engineering of wide-range antibacterial toxins.

WHIX is predicted to comprise a bipartite sequence that folds into a distinct domain containing a deep cavity. Based on our experimental findings and structure predictions, we propose that this cavity caps the C-terminal tail of a secreted spike component (e.g., VgrG), constituting a unique loading mechanism of T6SS effectors onto the tube-spike complex. In future work, we will determine the structure of WHIX domains and their mechanism of loading onto the secreted T6SS tube-spike. In addition, the role and identity of sequences separating the two regions of the WHIX domain remain to be investigated.

Interestingly, WHIX domains appear to specifically reside within effectors predicted to target the peptidoglycan, as indicated by the annotation of domains fused to WHIX. Such specificity of toxic domains to a cellular target was previously reported for T6SS effectors containing a FIX domain, although FIX domains are exclusively fused to toxic domains that target the bacterial cytoplasm (e.g., nucleases) [23]. Such target specificity implies that WHIX may play a role in the peptidoglycan-targeting activity of the effector, although WHIX itself was not toxic when ectopically expressed in the *E. coli* periplasm. Even though we showed that WHIX is sufficient for T6SS-mediated secretion, we cannot rule out the possibility that it also plays an active role, perhaps after deployment of the effector within the recipient periplasm.

We also characterize the *A. jandaei* T6SS in this work. We demonstrate that this T6SS is active under a wide range of temperatures and medium salinity in which it mediates antibacterial activity using at least four effectors. Although three of these effectors are encoded adjacent to secreted core components of the T6SS, either within the main T6SS gene cluster or within T6SS auxiliary operons, the fourth effector is orphan. We demonstrate that this effector, containing a DUF3289 domain recently predicted to be similar to colicin M [56,65], is part of a *bona fide* T6SS effector and immunity pair.

In conclusion, we describe a class of polymorphic T6SS cargo effectors defined by a unique secretion domain—WHIX. The presence of WHIX in double-edged sword effectors reveals a new mechanism used by bacteria to diversify their T6SS toxic effector payload. Our finding that WHIX effectors appear to predominantly target the peptidoglycan raises questions regarding the possibility of T6SS secretion domains also playing an active role in the activity of the effectors after deployment inside the recipient cell.

## Materials and Methods

### Strains and Media

For a complete list of strains used in this study, see Table S2. *Aeromonas jandaei* DSM 7311 and its derivative strains were grown in lysogeny broth (LB, 1% [wt/vol] tryptone, 0.5% [wt/vol] yeast extract, and 1% [wt/vol] NaCl) or LB agar (1.5% [w/v]) plates at 30ºC. *Escherichia coli* strains were grown in 2xYT broth (1.6% [wt/vol] tryptone, 1% [wt/vol] yeast extract, and 0.5% [wt/vol] NaCl) at 37°C, or at 30°C when harboring effector-expressing plasmids. In cases where *A. jandaei* or *E. coli* contain a plasmid, the media were supplemented with chloramphenicol (10 μg/mL) or kanamycin (30 μg/mL) in order to maintain the plasmid. To repress effector expression from the P*bad* promoter in *E. coli*, 0.4% D-glucose (wt/vol) was added to the media. To induce expression from the P*bad* promoter, L-arabinose (0.005%–0.1% [wt/vol]) was added to the media, as indicated.

### Plasmids Construction

For a complete list of plasmids used in this study, see Table S3. For expression in *A. jandaei* or *E. coli*, the coding sequence (CDS) of TssB (WP_033114640.1), Awe1 (WP_082035413.1) or its indicated truncations, Awi1U (WP_042029887.1), Awi1D (WP_156128651.1), DUF3289 (WP_042032936.1), the immunity protein encoded downstream of DUF3289 (encoded between position 2,059,343 and 2,059,057 in genome NZ_CP149571.1), and VgrG4 (WP_198493475.1) were PCR amplified from *A. jandaei* DSM 7311 genomic DNA; The CDS of superfolder GFP and GST were amplified from available plasmids. Amplicons were inserted into the multiple cloning site (MCS) of pBAD^K^/*Myc*-His, pBAD33.1 (Addgene), or their derivatives using the Gibson assembly method [66]. The resulting plasmids were introduced into *E. coli* DH5α (λ-pir) by electroporation, and into *A. jandaei* via conjugation. Trans-conjugants were selected on LB agar plates supplemented with the appropriate antibiotics. Since *A. jandaei* DSM 7311 is naturally resistant to ampicillin, it was added to the selection plates (50 μg/mL).

### Construction of Deletion Strains

To construct deletion strains, 1 kb sequences upstream and downstream of each gene or region to be deleted were cloned into the MCS of pDM4, a Cm^R^OriR6K suicide plasmid [67]. The pDM4 constructs were transformed into *E. coli* DH5α (λ-pir) by electroporation, and then transferred into *A. jandaei* strains via conjugation. Trans-conjugants were selected on agar plates supplemented with chloramphenicol (10 μg/mL) and ampicillin (50 μg/mL). The resulting trans-conjugants were grown on LB agar plates containing 10% (wt/vol) sucrose to select for the loss of the *sacB*-containing plasmid. Deletions were confirmed by PCR.

### Aeromonas Secretion Assays

For secretion of Hcp, Awe1 and its derivatives, VgrG4, and WP_042032936, *A. jandaei* strains were grown overnight in LB; appropriate antibiotics were included when required to maintain an expression plasmid. Bacterial cultures were normalized to OD_600_ = 0.5 in 5 mL LB supplemented with appropriate antibiotics and 0.005%-0.1% (wt/vol) L-arabinose as indicated when expression from an arabinose-inducible plasmid was required. Bacterial cultures were incubated with constant shaking (220 rpm) for 3 hours at 30ºC. After 3 hours, 0.5 OD_600_ units were collected for expression fractions (cells). Cell pellets were resuspended in (2x) Tris-Glycine SDS Sample Buffer (Novex, Life Sciences) supplemented with 5% (vol/vol) β-mercaptoethanol. For secretion fraction (media), culture volumes equivalent to 10 OD_600_ units were filtered (0.22 µm), and proteins were precipitated using the deoxycholate and trichloroacetic acid method [68]. Protein pellets were washed twice with cold acetone, air-dried for 10 minutes, and then resuspended in 20 µL of 10 mM Tris–HCl pH = 8.0, followed by the addition of 20 µL of (2x) Tris-Glycine SDS Sample Buffer supplemented with 5% (vol/vol) β-mercaptoethanol. Next, samples from the cells and media fractions were incubated at 95°C for 10 or 5 minutes, respectively, and loaded onto TGX stain-free gels (Bio-Rad) for SDS-PAGE. Resolved proteins were transferred onto 0.2 µm nitrocellulose membranes using Trans-Blot Turbo Transfer (Bio-Rad). Membranes were probed with custom-made anti-Hcp antibodies (GenScript; polyclonal antibodies raised in rabbits against the peptide DPQSGQPAGQRVHKC), anti-c-Myc antibodies (Santa Cruz Biotechnology, sc-40, 9E10), anti-FLAG antibodies (Sigma-Aldrich, F1804), or anti-GST antibodies (Santa Cruz Biotechnology; sc-459) at a 1:1000 dilution. As a control for loading and lysis, membranes were also probed with Direct-Blot™ HRP anti-*E. coli* RNA Polymerase β Antibody (Bio-legend, clone 8RB13; referred to as α-RpoB) at a 1:40,000 dilution. Protein signals were visualized in a Fusion FX6 imaging system (Vilber Lourmat) using ECL.

### Comparative Proteomics Analyses

Supernatant samples were obtained as described in the “*Aeromonas* Secretion Assays” section from three biological replicates for each strain. Pelleted proteins from the supernatant fractions were kept in acetone and sent to the Smoler Proteomics Center at the Technion for subsequent mass spectrometry analyses.

#### Proteolysis and Mass Spectrometry Analysis

The samples were precipitated in 80% acetone overnight and washed 3 times with 80% acetone. The protein pellets were dissolved in 8.5 M urea and 400 mM ammonium bicarbonate. Protein concentrations were estimated using Bradford readings. The proteins were reduced with 10 mM DTT (60°C for 30 min), modified with 40 mM iodoacetamide in 100 mM ammonium bicarbonate (at room temperature for 30 min in the dark), and digested overnight at 37°C in 1.5 M urea and 66 mM ammonium bicarbonate with modified trypsin (Promega) in a 1:50 (M/M) enzyme-to-substrate ratio. An additional trypsin digestion step was performed for 4 hours at 37°C in a 1:100 (M/M) enzyme-to-substrate ratio. The tryptic peptides were desalted using C18 tips (homemade stage tips), dried, and re-suspended in 2% ACN\H2O\ 0.1% Formic acid. The resulting peptides were analyzed by LC-MS/MS using a Q Exactive HF mass spectrometer (Thermo) fitted with a capillary HPLC (Evosep). The peptides were loaded onto a 15 cm, ID 150 µm, 1.9-micron (Batch no. E1121-3-24) column of Evosep. The peptides were eluted with the built-in Xcalibur 15 SPD (88 min) method. Mass spectrometry was performed in a positive mode using repetitively full MS scan (m/z 300–1500) followed by High energy Collision Dissociation (HCD) of the 20 most dominant ions selected from the full MS scan. A dynamic exclusion list was enabled with an exclusion duration of 20 sec.

#### Mass spectrometry data analysis

The MaxQuant software 2.1.1.0 [69] was used for peak picking and identification using the Andromeda search engine, searching against a locally annotated genome of *Aeromonas jandaei* DSM 7311, available in File S2, with mass tolerance of 6 ppm for the precursor masses and 20 ppm for the fragment ions. Oxidation on methionine and protein N-terminus acetylation were accepted as variable modifications and carbamidomethyl on cysteine was accepted as static modifications. The minimal peptide length was set to 7 amino acids and a maximum of two miscleavages was allowed. Peptide- and protein-level false discovery rates (FDRs) were filtered to 1% using the target-decoy strategy. The data was quantified by label-free analysis.

Protein tables were filtered to eliminate the identifications from the reverse database and common contaminants. Statistical analysis of the identification and quantization results was done using the Perseus 1.6.7.0 software [70] (Mathias Mann’s group).

### Bacterial Competition Assays

Bacterial competitions were performed as described previously [71]. Briefly, attacker and prey strains were grown overnight in appropriate media. Bacterial cultures were normalized to an OD_600_ = 0.5, and then mixed at a 4∶1 (attacker:prey) ratio in triplicate. The mixtures were spotted onto LB agar plates and incubated for 4 hours at 30°C. The plates were supplemented with 0.05%-0.1% (wt/vol) L-arabinose, as indicated, when expression from an arabinose-inducible plasmid was required. The viability of the prey strain was determined as colony-forming units growing on selective plates at the 0 and 4-hour time points.

### Toxicity Assays in E. coli

To examine the toxic effects of Awe1 and its derivatives, *E. coli* MG1655 strains carrying arabinose-inducible expression plasmids encoding the indicated proteins were grown overnight in 2xYT media supplemented with kanamycin (30 μg/mL) and 0.04% (wt/vol) D-glucose (to repress leaky expression from the *Pbad* promoter). Bacterial cultures were washed twice (to remove residual glucose) and normalized to OD_600_ = 0.01 in 2xYT media supplemented with kanamycin (30 μg/mL). From each sample, 200 µL were transferred into 96-well plates in quadruplicate. Cultures were grown at 30°C in a BioTek SYNERGY H1 microplate reader with constant shaking (205 cpm). After two ours, L-arabinose was added to each well to a final concentration of 0.05% (wt/vol), to induce expression from the plasmids. OD_600_ readings were taken every 10 minutes; the data were visualized using GraphPad PRISM.

### Protein Expression in E. coli

To examine the toxic effects of Awe1 and its derivatives, *E. coli* MG1655 strains carrying arabinose-inducible expression plasmids encoding the indicated proteins were grown overnight in 2xYT media supplemented with kanamycin (30 μg/mL) and 0.04% (wt/vol) D-glucose (to repress leaky expression from the *Pbad* promoter). Bacterial cultures were washed twice (to remove residual glucose) and normalized to OD_600_ = 0.5 in 3 mL 2xYT media containing kanamycin (30 μg/mL). Bacterial cultures were incubated with agitation at 30°C for 1.5 hours, and then L-arabinose was added to a final concentration of 0.05% (wt/vol) to induce protein expression. Bacterial cultures were grown for 1.5 additional hours, and then 0.5 OD_600_ units were collected, and pellets were resuspended in (2x) Tris-Glycine SDS Sample Buffer (Novex, Life Sciences) supplemented with 5% (vol/vol) β-mercaptoethanol. Next, samples were incubated at 95°C for 10 and loaded onto TGX stain-free gels (Bio-Rad) for SDS-PAGE. Transfer onto 0.2 µm nitrocellulose membranes was performed using Trans-Blot Turbo Transfer (Bio-Rad). Membranes were probed with anti-c-Myc antibodies (Santa Cruz Biotechnology, sc-40, 9E10) at a 1:1000 dilution. Protein signals were visualized in a Fusion FX6 imaging system (Vilber Lourmat) using ECL.

### Identifying WHIX Domain-Containing Proteins

PSI-BLAST was employed to construct the position-specific scoring matrix (PSSM) of WHIX. Five iterations of PSI-BLAST were performed against the RefSeq protein database using amino acids 1-477 of WP_005373349.1 (from *V. alginolyticus* 12G01). A maximum of 500 hits with an expect value threshold of 10^−6^ and 70% query coverage were used in each iteration.

A local database containing the RefSeq bacterial nucleotide and protein sequences was built (last updated on August 21, 2023). RPS-BLAST was used to identify WHIX domain-containing proteins in the local database. The results were filtered using an expect value threshold of 10^−6^ and a 70% overall coverage of the WHIX domain. In cases where more than one hit was identified in a protein accession, the boundaries of the WHIX domain in the protein accession were determined based on the starting position of the first hit and the ending position of the last hit.

Analysis of the genomic neighborhoods of WHIX domain-containing proteins (Dataset S1) was performed as described previously [72,73]. Unique protein accessions located at the ends of the contigs were removed. To avoid duplications, protein accessions appearing in the same genome in more than one genomic accession were removed if the same downstream protein existed at the same distance from the protein. Identification of the T6SS core components [6] in WHIX domain-containing genomes (Dataset S2) was performed as previously described [23].

Genomes encoding at least 9 out of the 11 T6SS core components were regarded as harboring T6SS (T6SS^+^). Protein accessions were analyzed using the NCBI Conserved Domain Database [58], SignalP v5.0 [74], and Phobius v1.01 [57].

### Illustration of Conserved Residues in WHIX Domains

WHIX domain sequences were aligned using Clustal-Omega v1.2.4 [75]. Aligned columns not found in the WHIX domain of WP_005373349.1, the protein accession that was used to generate the PSSM of WHIX, were discarded. The conserved WHIX domain residues were illustrated using the WebLogo 3 server [76] (https://weblogo.threeplusone.com).

### Constructing a Phylogenetic Tree of WHIX Domain-Containing Proteins

Q-TREE v2.2.2.6 [77] was employed to construct a Maximum-Likelihood phylogenetic tree of WHIX domain-containing proteins. The multiple sequence alignment of the WHIX domain, generated using Clustal-Omega, was used as input. 4,433 sequences with 2,484 amino acid sites were analyzed (360 invariant sites, 1,881 parsimony informative sites, and 2,305 distinct site patterns). ModelFinder [78] was used to test various protein models. The best-fit model, JTT+I+G4, was chosen according to the Bayesian Information Criterion (BIC). The tree was visualized using iTOL [79] (https://itol.embl.de/).

### Analyses of Domains Fused to WHIX

Domains fused to the C-terminal end of WHIX were identified according to annotations in the NCBI CDD [58]. To identify N-terminal domains in subclass II WHIX effectors (> 99 amino acids long), we clustered the N-terminal sequences in two dimensions using the CLANS application [80,81]. Activities or domains in each cluster were identified by analyzing at least two representative sequences from each cluster using the NCBI CDD [58], followed by analysis using HHpred [80] if no domain was apparent in CDD.

### Protein Structure Predictions

The structure prediction of WP_005373349.1, Awe1, and Awe1 in complex with AwiU and AwiD were carried out using AlphaFold 3 [46] (https://golgi.sandbox.google.com/). The best model was selected and visualized using ChimeraX [82] version 1.7.1 together with its Predicted Aligned Error (PAE) plot.

The structure prediction of the complex formed by monomeric Awe1 and trimeric VgrG4 was carried out using the AlphaFold-Multimer 2 [83,84]with AlphaFold Protein Structure Database (released on July 2022). The prediction process involved generating multiple models, with the best model selected based on the lowest PAE values. The final model and PAE plots were visualized using ChimeraX version 1.5. The specific interactions between Awe1 and VgrG4 were further analyzed by examining the predicted interfaces. Residues of Awe1 and VgrG predicted to interact were identified and highlighted in green and orange, respectively. All computations were performed on an in-house computer with an NVIDIA GeForce RTX 3080 GPU and 250 GB of RAM.

### Whole Genome Sequencing

*Aeromonas jandaei* DSM 7311 was purchased from DSMZ (https://www.dsmz.de/). Genomic DNA was isolated using the Presto mini gDNA Bacteria kit (Geneaid), and DNA was sent to Plasmidsaurus (https://www.plasmidsaurus.com/) for their bacterial genome sequencing service using [Oxford Nanopore Technologies (ONT)](https://nanoporetech.com/) long-read sequencing technology. Briefly, the process included: (i) construction of an amplification-free long read sequencing library using v14 library prep chemistry, (ii) sequencing the library using R10.4.1 flow cells, (iii) assembling the genome using the Plasmidsaurus standard pipeline, and (iv) genome annotation using Bakta [85].

The genome sequence is available as NCBI RefSeq assembly GCF_037890695.1; Chromosome RefSeq accession NZ_CP149571.1.

## Data Availability

The mass spectrometry proteomics data have been deposited in the ProteomeXchange Consortium via the PRIDE [86] partner repository with the dataset identifier PXD054809. The data can be downloaded via https://ftp.pride.ebi.ac.uk/pride/data/archive/2024/08/PXD054809.

The whole genome sequence of *A. jandaei* DSM 7311 is available as NCBI RefSeq assembly GCF_037890695.1; Chromosome RefSeq accession NZ_CP149571.1.

The AlphaFold-produced structure models used in this work are available as supplementary files.

## Acknowledgments

This project received funding from the Israel Science Foundation (ISF grant number 1362/21 to DS and EB). We thank the Smoler Proteomics Center at the Technion for performing and analyzing the mass spectrometry data.

## Author contributions

Conceptualization: EB and DS; Formal Analysis: CMF, DAJ, EB, and DS; Funding Acquisition: EB, and DS; Investigation: CMF, KK, VR, JAA, EB, and DS; Methodology: CMF, DAJ, and EB; Resources: DAJ and EB; Supervision: DS; Writing – Original Draft Preparation: DS; Writing – Review and Editing: CMF, KK, VR, JAA, DAJ, and EB.

