## Supplementary material for "WHIX is a T6SS secretion domain found in polymorphic double-edged sword effectors": Supplemenal Information

Supplementary Figures S1-S5

Supplementary Tables S1-S3

Supplementary Dataset S1-S2 (captions)

Supplementary Files S1-S7 (captions)

Supplementary References

### Supplementary Figures

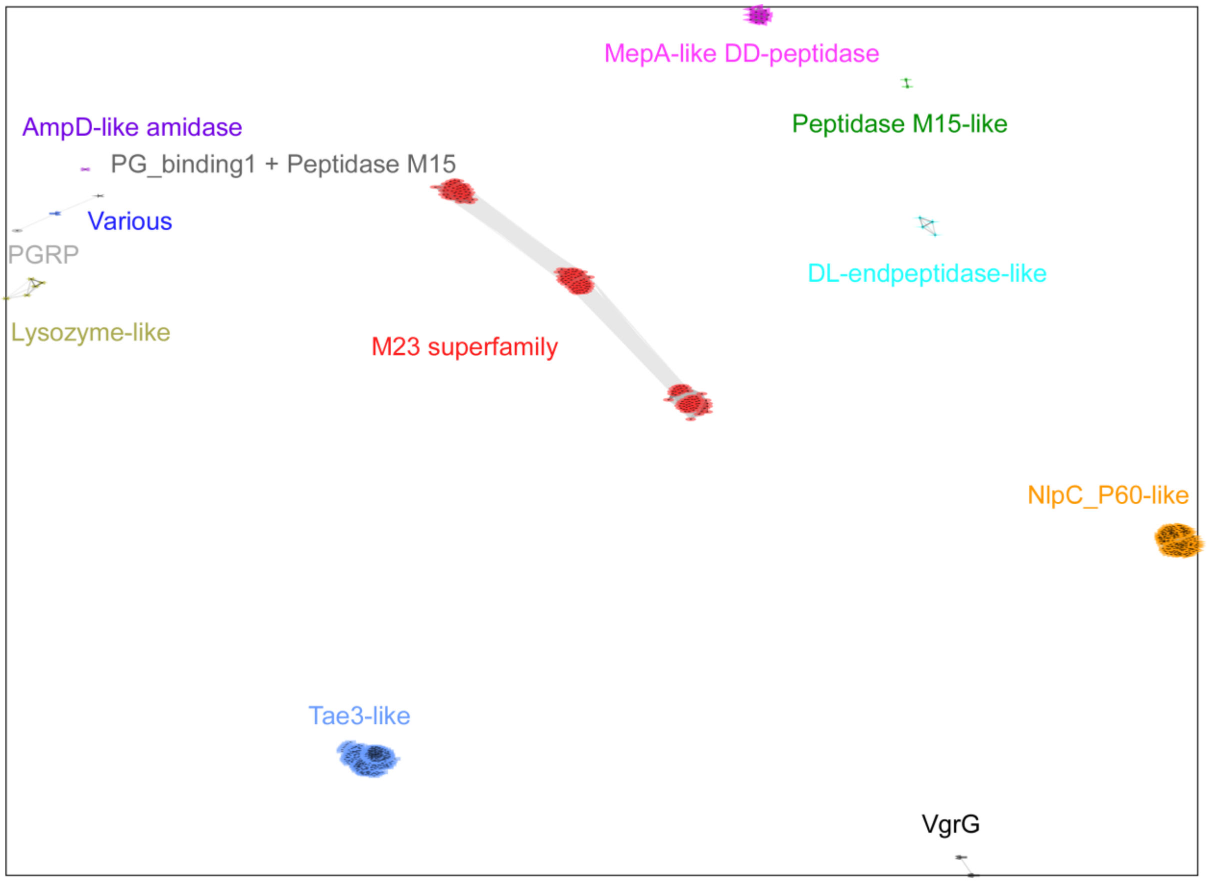


**Fig S1: N-terminal domains in subclass II WHIX effectors comprise diverse peptidoglycan-targeting enzymes.** Sequences N-terminal to WHIX domains in subclass II WHIX effectors were clustered in two dimensions based on all-against-all sequence similarity, using the CLANS application, with nodes representing unique sequences and connecting lines representing the distances between sequences. The predicted activities or domains identified in each cluster are denoted and color-coded according to the nodes.

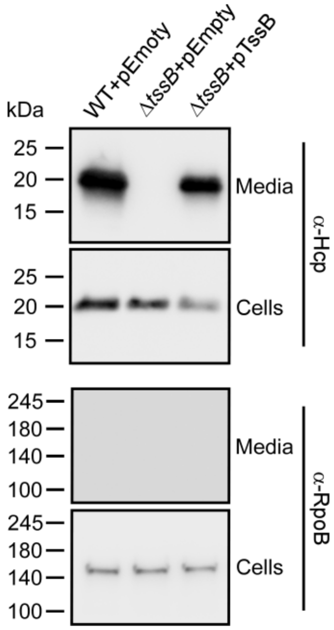


**Fig S2: *A. jandaei* DSM 7311 has a functional T6SS.** Expression (cells) and secretion (media) of Hcp from wild-type (WT) *A. jandaei* DSM 7311 and a T6SS$^-$ mutant strain (Δ*tssB*), containing either an empty plasmid (pEmpty) or a plasmid for the arabinose-inducible expression of *tssB* (pTssB), grown for 3 hours at 30°C in LB media supplemented with chloramphenicol and 0.1% L-arabinose. RNA polymerase beta subunit (RpoB) was used as a loading and lysis control. Results from a representative experiment out of at least three independent experiments are shown.

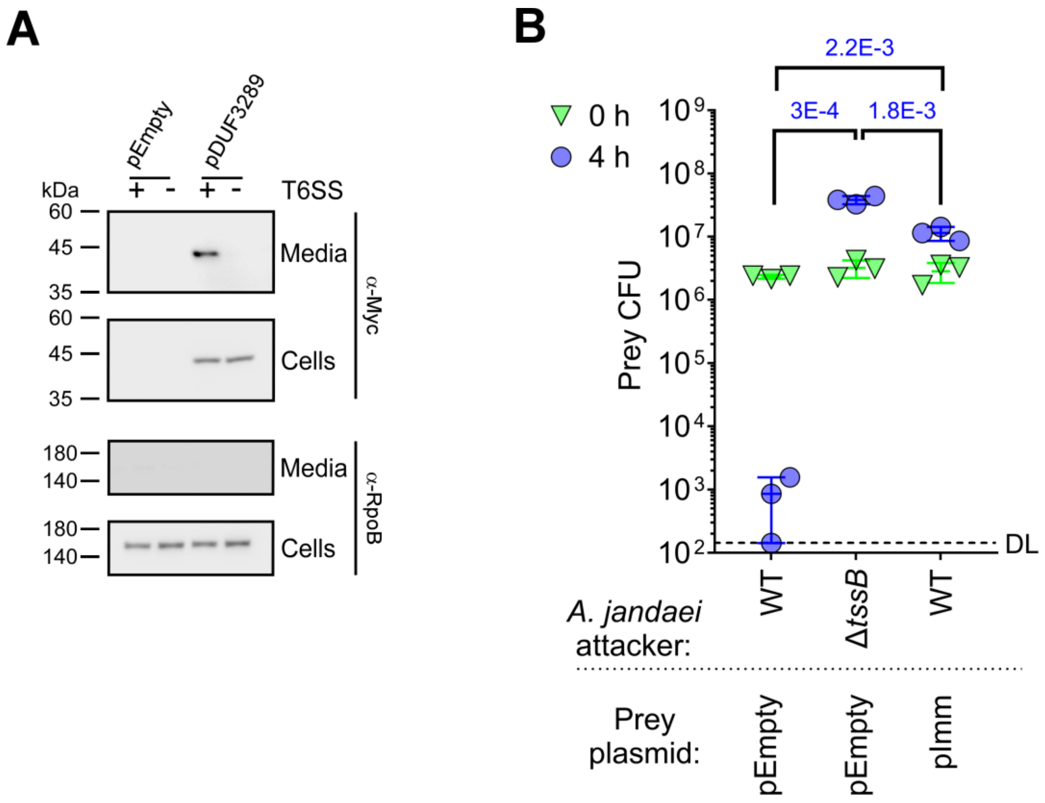


**Fig S3: WP_042032936.1 is an antibacterial T6SS effector. (A)** Expression (cells) and secretion (media) of FLAG-tagged WP_042032936.1 expressed from an arabinose-inducible plasmid (pDUF3289) in wild-type (WT) *A. jandaei* DSM 7311 and a T6SS$^-$ mutant strain (Δ*tssB*) grown for 3 hours at 30°C in LB media supplemented with kanamycin and 0.05% L-arabinose. RNA polymerase beta subunit (RpoB) was used as a loading and lysis control. Results from a representative experiment out of at least three independent experiments are shown. **(B)** Viability counts (CFU) of *A. jandaei* DSM 7311 prey strains in which the genes encoding WP_042032936.1 and its predicted cognate immunity protein were deleted, containing an empty plasmid (pEmpty) or a plasmid for the arabinose-inducible expression of the predicted immunity protein (pImm), before (0 h) and after (4 h) co-incubation with the indicated *A. jandaei* DSM 7311 attacker strains on LB plates supplemented with 0.05% (wt/vol) L-arabinose at 30°C. The statistical significance between samples at the 4 h time point was calculated using an unpaired, two-tailed Student's *t* test; ns, no significant difference ($P > 0.05$); WT, wild-type; DL, the assay's detection limit. Data are shown as the mean ± SD; $n = 3$. The data shown are a representative experiment out of at least three independent experiments.

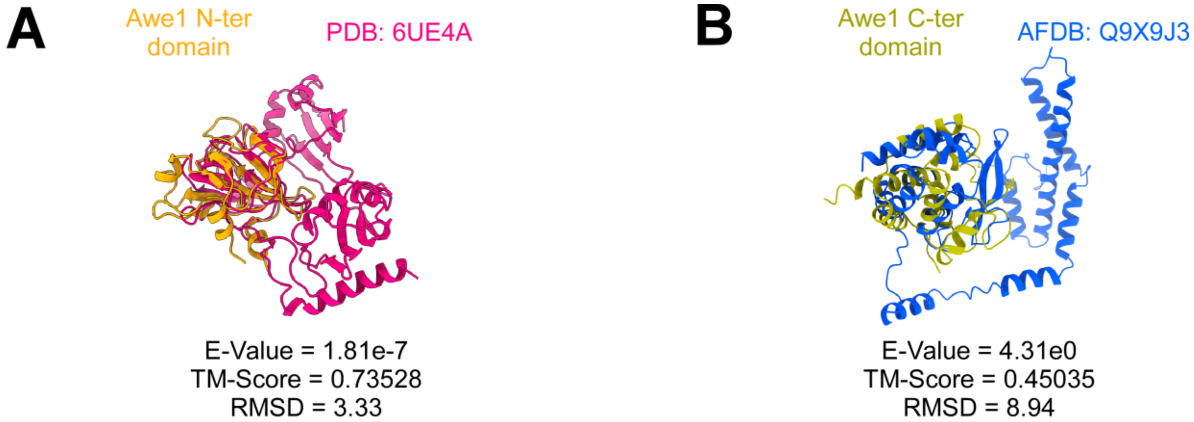


**Fig S4. Awe1 N- and C-terminal domains are similar to peptidoglycan-degrading enzymes. (A)** Superimposed AlphaFold3 structure prediction of the N-terminal (N-ter) Awe1 domain (amino acids 1-147; colored orange) with *V. cholerae* ShyA (PDB: 6UE4A; colored fiusha). **(B)** Superimposed AlphaFold3 structure prediction of the C-terminal (C-ter) Awe1 domain (amino acids 700-862; colored beige) with the AlphaFold structure prediction of *V. parahaemolyticus* FlgJ (AlphaFold database [AFDB]: Q9X9J3; colored blue).

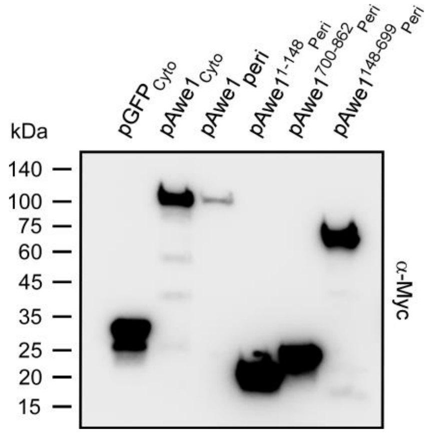


**Fig S5. Awe1 expression in *E. coli*.** The expression of the indicated C-terminally Myc-tagged superfolder GFP and Awe1 forms expressed in the cytoplasm (Cyto) or periplasm (Peri) of *E. coli* MG1655 from an arabinose-inducible plasmid.

### Supplementary Tables

**Table S1. Domains and activities identified at N-termini of subclass II WHIX effectors.**

| Number of unique proteins | Predicted domain | Example accession number |
|---|---|---|
| 328 | Tae3-like [a] | WP_282812471.1 |
| 222 | NlpC_P60-like [b] | WP_205350834.1 |
| 162 | M23 superfamily [b] | WP_069143508.1 |
| 13 | MepA-like DD-peptidase [a] | WP_158695274.1 |
| 5 | Lysozyme-like [b] | WP_281319639.1 |
| 4 | DL-endopeptidase-like [a] | WP_267540890.1 |
| 2 | Peptidase M15-like [b] | WP_206728780.1 |
| 2 | VgrG [b] | WP_062088695.1 |
| 1 | AmpD-like amidase [b] | WP_044238255.1 |
| 1 | PGRP [b] | WP_181197892.1 |
| 1 | PG_binding1 + Peptidase M15 [a,b] | WP_266216180.1 |
| 1 | Various [b] | WP_284721687.1 |

[a] Identified using HHpred analyses, [b] Identified using the NCBI CDD

**Table S2. Bacterial strains used in this study.**

| Strain name | Genotype | Comments | Source |
|---|---|---|---|
| *Aeromonas jandaei* DSM 7311 | Wild-type | Used in competition assays, secretion assays, and for generating deletion strains. The strain is also named ATCC 49568 and CECT 4228 | DSMZ collection |
| *Aeromonas jandaei* Δ*tssB* | DSM 7311 Δ*we862_rs13035* | Used in competition and secretion assays | [1] |
| *Aeromonas jandaei* Δ*awe1* | DSM 7311 Δ*we862_rs20670* | Used in competition and secretion assays | This study |
| *Aeromonas jandaei* Δ*awe1*/Δ*tssB* | DSM 7311 Δ*we862_rs13035* / Δ*we862_rs20670* | Used in competition and secretion assays | This study |
| *Aeromonas jandaei* ΔI-E-I | DSM 7311 Δ*we862_rs20675*-*we862_rs20665* | Used in competition assays | This study |
| *Aeromonas jandaei* Δ*vgrG4* | DSM 7311 Δ*we862_20680* | Used in competition and secretion assays | This study |
| *Aeromonas jandaei* Δ*vgrG4*/Δ*tssB* | DSM 7311 Δ*we862_20680* / Δ*we862_rs13035* | Used in competition and secretion assays | This study |
| *Aeromonas jandaei* Δ*duf3289*-Imm | DSM 7311 Δ*we862_rs09995*-downstream immunity (not annotated on NCBI) | Used in competition assays | This study |
| *Aeromonas jandaei* *Aj*$^{effectorless}$ | DSM 7311 Δ*we862_rs20670* / Δ*we862_rs16925* / Δ*we862_rs09995* / Δ*we862_rs13125* (nucleotides encoding amino acids 779-1509) | Used in competition assays | This study |
| *Escherichia coli* DH5α (λ-pir) | K-12 derivative laboratory strain containing λ-pir | Used for plasmid maintenance and cloning | Obtained from Eric V. Stabb |
| *Escherichia coli* BL21 (DE3) | Laboratory strain | Used as a prey in competition assays | Lab stocks |
| *Escherichia coli* MG1655 | Wild-type | Used in competition and toxicity assay | Lab stocks |

**Table S3. Plasmids used in this study.**

| Plasmid name | Description | Purpose | Source |
|---|---|---|---|
| pBAD$^K$/*Myc*-His | pBR322 ori-containing plasmid harboring a Kan$^R$ cassette, *araC*, and an MCS following a P*bad* promoter. A Myc-His tag is encoded at the 5' end of the MCS. | Used as an empty plasmid control | [2] |
| pAwe1 (also named pAwe1$_{Cyto}$) | pBAD$^K$/*Myc*-His plasmid containing *awe1*, in-frame with a C-terminal Myc-His tag | Used for arabinose-inducible expression of Awe1 | This study |
| pAwe1$^{1-699}$ | pBAD$^K$/*Myc*-His plasmid containing the region in *awe1* encoding Awe1$^{1-699}$, in-frame with a C-terminal Myc-His tag | Used for arabinose-inducible expression of Awe1$^{1-699}$ | This study |
| pAwe1$^{148-862}$ | pBAD$^K$/*Myc*-His plasmid containing the region in *awe1* encoding Awe1$^{148-862}$, in-frame with a C-terminal Myc-His tag | Used for arabinose-inducible expression of Awe1$^{148-862}$ | This study |
| pAwe1$^{148-699}$ | pBAD$^K$/*Myc*-His plasmid containing the region in *awe1* encoding Awe1$^{148-699}$, in-frame with a C-terminal Myc-His tag | Used for arabinose-inducible expression of Awe1$^{148-699}$ | This study |
| pGEX4T-1 | Lactose analog isopropyl β-D-thiogalactoside (IPTG)-inducible pGEX plasmid harboring an AmpR cassette | Used as a template for amplifying GST | GE Healthcare |
| pGST | pBAD$^K$/*Myc*-His plasmid encoding GST (untagged) | Used for arabinose-inducible expression of GST | This study |
| pGST-Awe1$^{148-862}$ | pBAD$^K$/*Myc*-His plasmid encoding an N-terminal GST followed by a AAAGGG linker and Awe1$^{148-862}$, in-frame with a C-terminal Myc-His tag | Used for arabinose-inducible expression of GST- Awe1$^{148-862}$ | This study |
| pGST-Awe1$^{148-699}$ | pBAD$^K$/*Myc*-His plasmid encoding an N-terminal GST followed by a AAAGGG linker and Awe1$^{148-699}$, in-frame with a C-terminal Myc-His tag | Used for arabinose-inducible expression of GST- Awe1$^{148-699}$ | This study |
| pGST-Awe1$^{148-699}$-GFP | pBAD$^K$/*Myc*-His plasmid encoding an N-terminal GST followed by a AAAGGG linker, Awe1$^{148-699}$, another AAAGGG linker, and sfGFP, in-frame with a C-terminal Myc-His tag | Used for arabinose-inducible expression of GST- Awe1$^{148-699}$-GFP | This study |

| | | | |
|---|---|---|---|
| pGFP | pBAD$^{K}$/*Myc*-His plasmid encoding sfGFP, in-frame with a C-terminal Myc-His tag | Used for arabinose-inducible expression of GFP | This study |
| psfGFP | pBAD33.1$^{F}$ plasmid containing the CDS of sfGFP in-frame with a C-terminal FLAG tag | Used as template to amplify sfGFP | [3] |
| pAwe1$^{1-699}$-GFP | pBAD$^{K}$/*Myc*-His plasmid encoding an Awe1$^{1-699}$, a AAAGGG linker, and sfGFP, in-frame with a C-terminal Myc-His tag | Used for arabinose-inducible expression of Awe1$^{1-699}$-GFP | This study |
| pAwe1$^{148-699}$-GFP | pBAD$^{K}$/*Myc*-His plasmid encoding an Awe1$^{148-699}$, a AAAGGG linker, and sfGFP, in-frame with a C-terminal Myc-His tag | Used for arabinose-inducible expression of Awe1$^{148-699}$-GFP | This study |
| pDUF3289 | pBAD$^{K}$/*Myc*-His plasmid containing the CDS of WP_042032936.1, in-frame with a C-terminal Myc-His tag | Used for arabinose-inducible expression of WP_042032936.1 | This study |
| pPER5 | pBADK/Myc-His with a PelB signal peptide inserted at the 5' end of the MCS | Used for arabinose-inducible expression of proteins targeted to the periplasm | [4] |
| pAwe1$_{peri}$ | pPER5 plasmid containing the CDS of Awe1, in-frame with an N-terminal PelB signal peptide and a C-terminal Myc-His tag | Used for arabinose-inducible expression of Awe1 in the periplasm | This study |
| pAwe1$^{1-147}_{peri}$ | pPER5 plasmid containing the CDS of Awe1$^{1-147}$, in-frame with an N-terminal PelB signal peptide and a C-terminal Myc-His tag | Used for arabinose-inducible expression of Awe1$^{1-147}$ in the periplasm | This study |
| pAwe1$^{700-862}_{peri}$ | pPER5 plasmid containing the CDS of Awe1$^{700-862}$, in-frame with an N-terminal PelB signal peptide and a C-terminal Myc-His tag | Used for arabinose-inducible expression of Awe1$^{700-862}$ in the periplasm | This study |
| pAwe1$^{148-699}_{peri}$ | pPER5 plasmid containing the CDS of Awe1$^{148-699}$, in-frame with an N-terminal PelB signal peptide and a C-terminal Myc-His tag | Used for arabinose-inducible expression of Awe1$^{148-699}$ in the periplasm | This study |
| pBAD33.1 | p15A ori-containing plasmid carrying a Cm$^{R}$ gene, *araC*, and an MCS following a P*bad* promoter | Used as an empty plasmid control | Addgene |

| | | | |
|---|---|---|---|
| pBAD33.1$^{F}$ | pBAD33.1 with a FLAG tag inserted at the 3' end of the MCS | Used for arabinose-inducible expression of proteins with a C-terminal FLAG tag | [5] |
| pBAD33.1$^{NF}$ | pBAD33.1 with a FLAG tag inserted at the 5' end of the MCS | Used for arabinose-inducible expression of proteins with an N-terminal FLAG tag | This study |
| pTssB | pBAD33.1$^{F}$ plasmid containing the CDS of TssB from *A. jandaei* DSM 7311 (untagged) | Used for arabinose-inducible expression of TssB | This study |
| pVgrG4 | pBAD33.1$^{NF}$ plasmid containing the CDS of VgrG4 from *A. jandaei* DSM 7311, in-frame with an N-terminal FLAG tag | Used for arabinose-inducible expression of VgrG4 in *A. jandaei* | This study |
| pVgrG4$^{1-680}$ | pBAD33.1$^{NF}$ plasmid encoding VgrG4$^{1-680}$ from *A. jandaei* DSM 7311, in-frame with an N-terminal FLAG tag | Used for arabinose-inducible expression of VgrG4$^{1-680}$ in *A. jandaei* | This study |
| pAwiU | pBAD33.1$^{F}$ plasmid containing the CDS of AwiU, in-frame with a C-terminal FLAG tag | Used for arabinose-inducible expression of AwiU | This study |
| pAwiD | pBAD33.1$^{F}$ plasmid containing the CDS of AwiD, in frame with a C-terminal FLAG tag | Used for arabinose-inducible expression of AwiD | This study |
| pAwiU+ pAwiD | pBAD33.1$^{F}$ plasmid containing the CDS of AwiU in the MCS, followed by another P*bad* promoter and the CDS of AwiD; Both AwiU and AwiD are in-frame with a C-terminal FLAG tag | Used for arabinose-inducible expression of AwiU and AwiD together | This study |
| pImm | pBAD33.1$^{F}$ plasmid containing the CDS of the DUF3289 immunity gene, in frame with a C-terminal FLAG tag | Used for arabinose-inducible expression of DUF3289 immunity protein | This study |
| pDM4 | a Cm$^{R}$ and oriVR6K-containing suicide vector | Used as a template for constructing plasmids for gene deletions | [6] |
| pDM4:*tssB* | pDM4 containing 1 kb upstream and 1 kb downstream of the gene encoding | Used to delete *tssB* in *A. jandaei* DSM 7311 | [1] |

| | | | |
|---|---|---|---|
| | TssB in *A. jandaei* DSM 7311 in its MCS | | |
| pDM4:awe1 | pDM4 containing 1 kb upstream and 1 kb downstream of the gene encoding Awe1 in *A. jandaei* DSM 7311 in its MCS | Used to delete *awe1* in *A. jandaei* DSM 7311 | This study |
| pDM4:*awiU-awe1-awiD* | pDM4 containing 1 kb upstream of the gene encoding AwiU and 1 kb downstream of the gene encoding AwiD in *A. jandaei* DSM 7311 in its MCS | Used to delete *awiU-awe1-awiD* in *A. jandaei* DSM 7311 | This study |
| pDM4:*tle1* | pDM4 containing 1 kb upstream and 1 kb downstream of the gene encoding Tle1 in *A. jandaei* DSM 7311 in its MCS | Used to delete *tle1* in *A. jandaei* DSM 7311 | This study |
| pDM4:*tseI* | pDM4 containing 1 kb upstream and 1 kb downstream of the region encoding amino acids 779-1509 of TseI in *A. jandaei* DSM 7311 in its MCS | Used to delete *tseI* in *A. jandaei* DSM 7311 | This study |
| pDM4:*duf3289-imm* | pDM4 containing 1 kb upstream of the gene encoding the DUF3289 effector and 1 kb downstream of the gene encoding its cognate immunity protein in *A. jandaei* DSM 7311 in its MCS | Used to delete *duf3289* and its downstream immunity gene in *A. jandaei* DSM 7311 | This study |
| pDM4:*vgrG4* | pDM4 containing 1 kb upstream and 1 kb downstream of the gene encoding VgrG4 in *A. jandaei* DSM 7311 in its MCS | Used to delete *vgrG4* in *A. jandaei* DSM 7311 | This study |

### Supplementary Dataset captions

**Dataset S1: A list of WHIX-containing proteins and their genomic neighborhood.**

**Dataset S2: Identification of T6SSs in genomes encoding WHIX-containing proteins.**

### Supplementary File captions

**File S1: AlphaFold3 prediction of WP_005373349.1 structure.**

**File S2: CLANS analysis of sequences found N-terminal to WHIX in subclass II WHIX effectors.**

**File S3: Comparative proteomics mass spectrometry data analysis and the *A. jandaei* DSM 7311 genome annotation used for it.**

**File S4: AlphaFold3 prediction of Awe1 structure.**

**File S5: AlphaFold3 prediction of Awe1 structure in complex with AwiU and AwiD.**

**File S6: Superimposition of the Awe1 N- and C-terminal domains with peptidoglycan-degrading enzymes.**

**File S7: AlphaFold prediction of Awe1 structure in complex with a VgrG4 trimer.**
